# Sustainable Edamame Production in an Artificial Light Plant Factory with Improved Yield and Quality

**DOI:** 10.1101/2025.05.01.651627

**Authors:** Tomoki Takano, Yu Wakabayashi, Soshi Wada, Toshio Sano, Saneyuki Kawabata, Wataru Yamori

**Affiliations:** Graduate School of Agricultural and Life Sciences, The University of Tokyo, Nishitokyo, Tokyo, Japan; Faculty of Biosciences, Department of Clinical Plant Science, Hosei University, Tokyo, Japan

## Abstract

A plant factory utilizing artificial light is an innovative agricultural model that enables efficient and sustainable crop production. However, its application to a diverse range of crops remains limited. Edamame, a highly nutritious legume, has gained global popularity, yet its long-term storage is challenging due to quality deterioration, restricting its market distribution to seasonal availability. In this study, we successfully cultivated edamame using three hydroponic systems— nutrient film technique (NFT), rockwool, and aeroponics (mist culture)—within an LED plant factory. Among these, NFT demonstrated the highest fresh seed yield, which was comparable to or exceeded that of conventional field cultivation. The high yield was attributed to enhanced pod formation and biomass production under controlled conditions. Furthermore, although free amino acid content was lower in NFT cultivation, NFT-grown edamame exhibited significantly higher total sugar and isoflavone contents than field-grown counterparts, suggesting superior eating quality and nutritional benefits. These results indicate that NFT hydroponics is a promising method for stable, high-yield edamame production with enhanced nutritional properties. This study provides a foundation for expanding edamame cultivation in controlled environments, contributing to year-round supply and its potential application in urban and space agriculture.

## Introduction

The world’s population is projected to reach 8.5 billion by 2030 and 9.7 billion by 2050^1^. At the same time, climate change is negatively impacting food production through long-term shifts in air temperature, drought, pest infestations and so on^2–4^. Approximately one-third of global crop production relies on pesticide applications, and crop yields would decline drastically without pesticide use. Meanwhile, improper pesticide use leads to serious environmental pollution^5^. In this context, the introduction of plant factories is required to transition away from conventional agriculture. A plant factory is an advanced facility that enables continuous and automated year-round crop production by artificially controlling environmental conditions such as light, temperature, humidity, carbon dioxide levels, and nutrient composition in the culture medium, offering great potential for maximizing yields in limited cultivation spaces^6^ while minimizing pesticide use through the maintenance of optimal growth conditions^7^.

As global population growth demands a further increase in overall food production, protein-energy malnutrition remains a serious issue in developing countries^8^, highlighting the need to enhance protein source production. Legumes contain more than 20% protein in the edible part, the cotyledons^9^. Soybean is one of the most economically significant legume crops, accounting for over a quarter of the global protein supply, including livestock feed^10^. Moreover, soybeans contain 52% linoleic acid among their fatty acids—an omega-6 unsaturated fatty acid known for its health benefits^11^. Additionally, soybean-derived isoflavones have also gained attention for their potential health benefits, as they are expected to lower the risk of cancer and ischemic heart disease^12–14^.

Edamame refers to soybeans harvested at the reproductive growth stages, when the seeds are still green and immature^15^. Edamame has long been cultivated and consumed in East Asia, particularly in Japan and China. In recent decades, it has gained considerable attention in the United States and other regions due to its high nutritional value and great taste^16,17^. Compared to mature soybeans, edamame cotyledons contain higher levels of free amino acids and sugars^18^. The free amino acids mainly consist of asparagine, glutamic acid, alanine, arginine, serine, and GABA, while the sugars are primarily composed of sucrose, fructose, and glucose^19–21^, all of which contribute to its characteristic flavor^20^. Additionally, edamame is rich in isoflavones such as daidzin, malonyl daidzin, glycitein, genistin, and malonyl genistin, which are also considered to influence its flavor profile^18^. Due to the necessity of harvesting and shipping in a fresh state, edamame has a short storage period. Previous studies have shown that nutritional components such as free amino acids, sugars, and ascorbic acid decline sharply within two days of storage at 20°C after harvesting^22,23^. To date, several methods—including storage at approximately 0°C, keeping the cotyledons attached to branches, and improving film packaging— have been proposed to maintain freshness, however, none provide a complete solution. Therefore, unlike mature soybeans or other fruiting vegetables, long-term storage of edamame remains challenging. The Japanese Ministry of Agriculture, Forestry and Fisheries (MAFF) reports that its market distribution is concentrated in the summer, when harvest volumes are relatively high under natural conditions, while reduced availability in the winter leads to increased market prices.

In plant factories, various cultivation systems are utilized, however, each method has its own advantages and disadvantages. Solid substrate culture systems, such as rockwool, provide greater buffering capacity, making them less susceptible to fluctuations in crop conditions, however, their high buffering capacity also limits precise control over the composition of the culture medium^24^. In contrast, hydroponic and mist culture systems do not use solid substrates. Hydroponics is characterized by low buffering capacity, as nutrient solutions are supplied directly to plants, allowing for more precise and rapid control of culture solution components. However, this system is more sensitive to fluctuations in crop conditions. Moreover, mist culture is a cultivation method in which the culture solution is sprayed onto roots that are exposed to the gaseous phase, enabling crops to grow in an oxygen-rich root environment^25,26^. Nevertheless, regulating the supply of nutrients and water in mist culture systems is technically challenging, as the roots are suspended in air^27^. Solid substrate culture systems are generally employed for fruiting vegetables with long growth periods, such as tomatoes, cucumbers, strawberries, and bell peppers, whereas, hydroponics is primarily used for cultivating leafy vegetables and herbs, which have shorter growth periods and can be produced in faster cultivation cycles^28^. The selection of an optimal cultivation system depends on multiple factors, including crop growth, production quality, profitability, labor requirements, technical complexity, and environmental impact^29^. Ultimately, the most suitable cultivation method varies depending on the specific characteristics and requirements of the plant being grown.

The objective of this study was to develop an optimal cultivation system for edamame in a plant factory. Specifically, we aimed to achieve successful cultivation in an artificial light-type plant factory, with a view to future applications in urban agriculture and barren areas such as desert regions. Three cultivation systems were investigated in this study: nutrient film technique (NFT), which is predominantly used for lettuce cultivation in artificial light-type plant factories; rockwool culture (ROC), which is widely used for tomato cultivation in sunlight-type plant factories; and mist culture (MIST), which is considered advantageous for its potential application in rhizobium symbiosis. The optimal system for edamame cultivation in artificial light-type plant factories was determined through a comprehensive evaluation of plant growth, seed yield, and seed nutritional components. Furthermore, its productivity and quality potential were evaluated by comparison with field cultivation (FIELD).

## Results

### Growth traits

Plant growth traits were analyzed across three hydroponic cultivation systems—nutrient film technique (NFT), rockwool culture (ROC), and mist culture (MIST)—as well as field cultivation (FIELD) (Fig. 1). The stem length, number of leaflets, and number of nodes during the vegetative stage were generally highest in NFT, followed by ROC and MIST for both cultivars, with some exceptions (Fig. S1). The dry matter of each organ at maturity was compared across cultivation systems, including FIELD (Fig. S2). For Sayane, the stem dry weight was highest in NFT, followed by FIELD, ROC, and MIST, whereas no significant differences were observed among cultivation systems for Ooyukimidori. The leaf dry weight was significantly higher in NFT for Sayane and in NFT and ROC for Ooyukimidori compared with the other cultivation systems. The pod dry weight in NFT and ROC was higher than in MIST and comparable to FIELD. Similarly, the seed dry weight was significantly higher in NFT and ROC compared with MIST for both cultivars. Additionally, in Sayane, NFT and ROC had significantly higher seed dry weight than FIELD, whereas no significant differences were found for Ooyukimidori. The total above-ground dry weight was significantly higher in NFT and ROC than in MIST for both cultivars, and for Sayane, it was also higher than in FIELD. These results indicate that NFT and ROC promoted superior plant growth throughout the entire growth period.

**Fig. 1.**
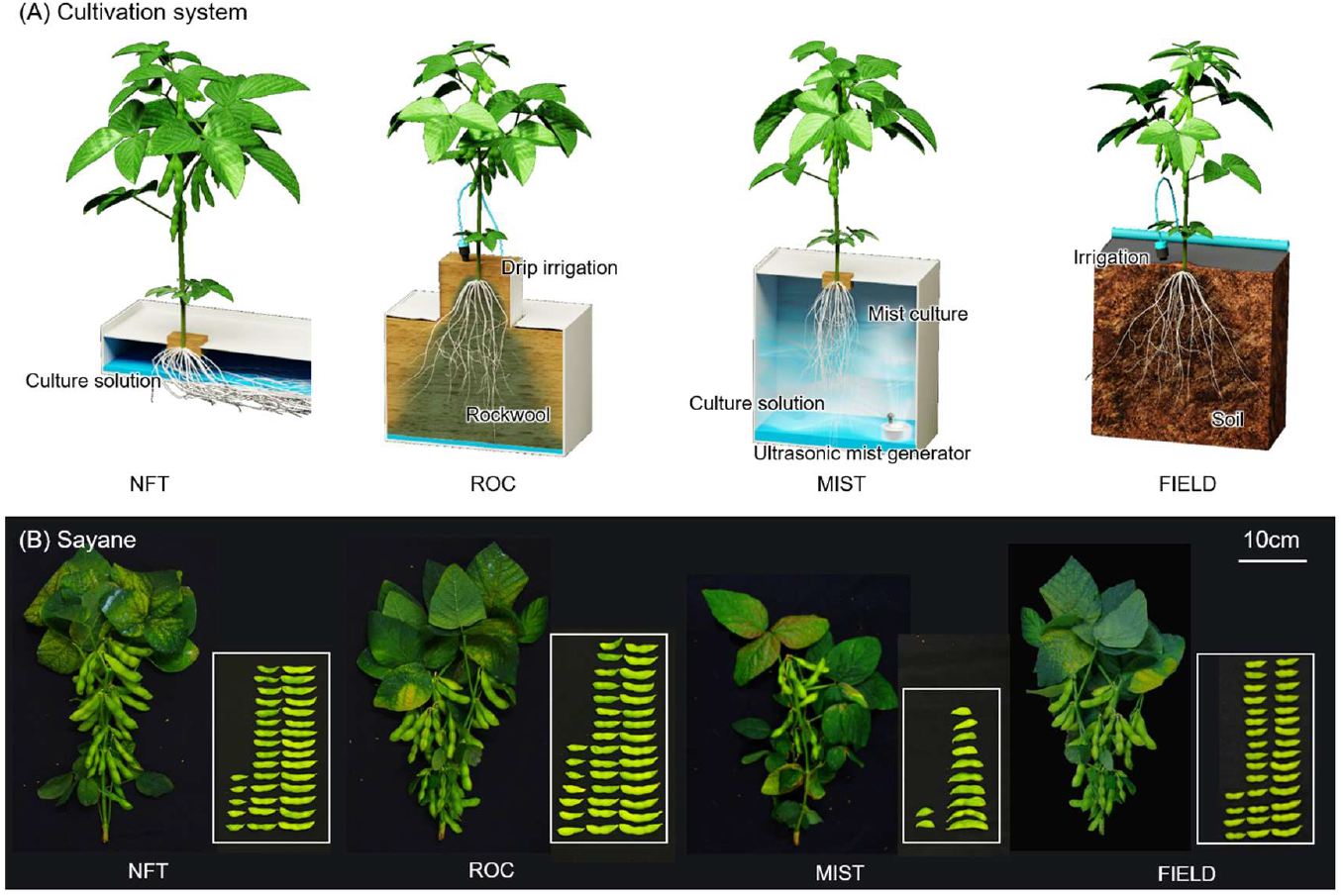
Cultivation systems in the present study. (A) Schematic diagram of each cultivation system. (B) Gross morphology of plants and pods at maturity.

### Yield-related traits

At 49 DAS, there were no significant differences in flower number among cultivation systems for either cultivar, although between 33 and 41 DAS, flower numbers were higher in ROC and NFT (Fig. S3). The pod number and seeds per pod were higher in NFT and ROC compared with MIST, while no significant differences were observed compared with FIELD for either cultivar (Fig. 2A). The fresh weight of a mature seed did not significantly differ among the plant factory cultivation systems for either cultivar. However, in Sayane, fresh seed weight was significantly higher in NFT and ROC compared with FIELD. Conversely, the moisture content of mature seeds was lower in NFT and ROC compared with MIST and FIELD for both cultivars. The percentage of mature seeds did not significantly differ among cultivation systems for either cultivar. The harvest index, calculated as the ratio of mature seed dry weight to total above-ground dry weight, was significantly higher in NFT and ROC than in MIST for both cultivars. In Sayane, NFT and ROC also showed a higher harvest index compared with FIELD. The fresh seed yield was significantly higher in NFT and ROC than in MIST for both cultivars. Furthermore, for Sayane, NFT yielded significantly more fresh seeds than FIELD, whereas for Ooyukimidori, NFT yielded a comparable amount to FIELD.

**Fig. 2.**
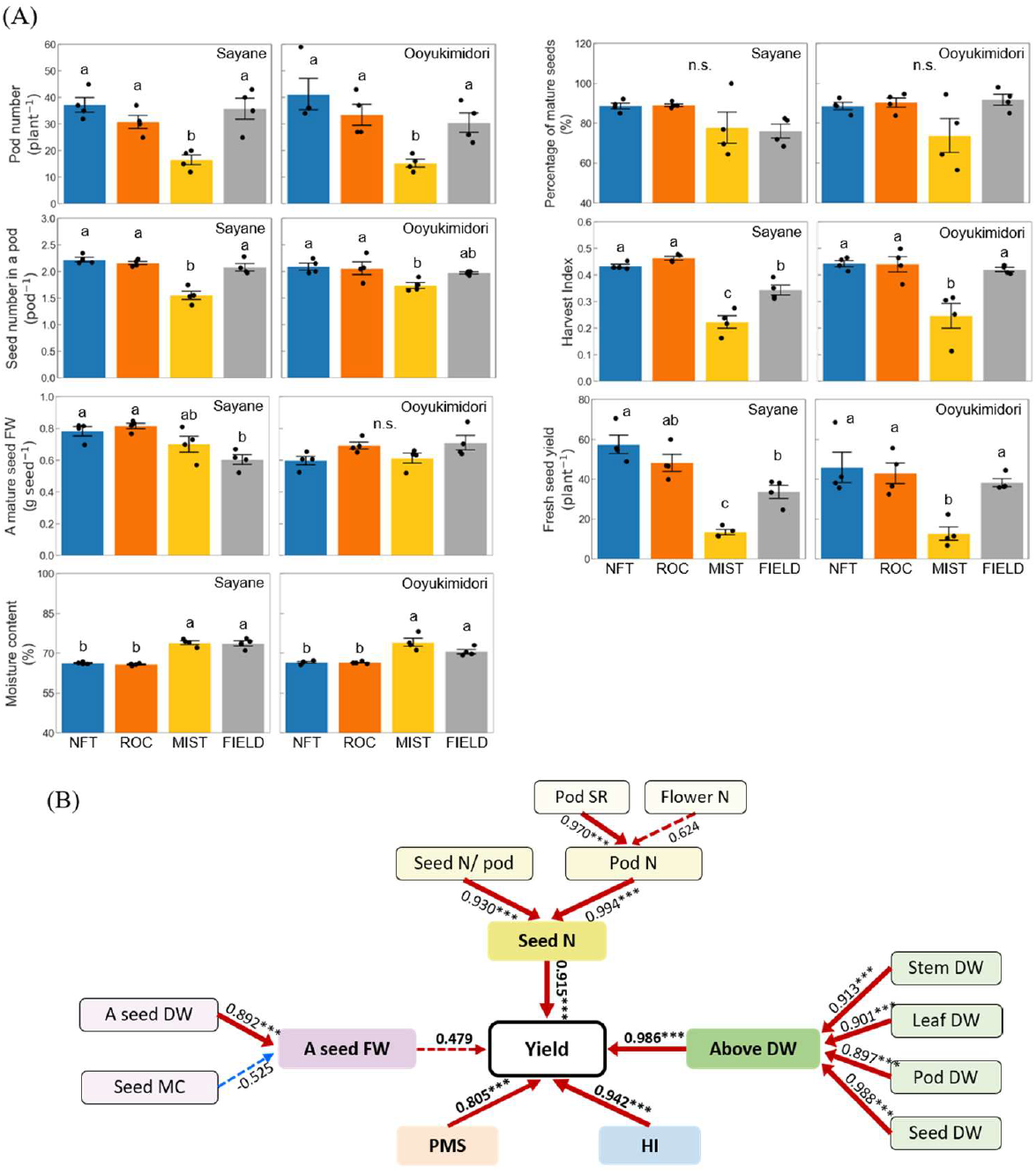
Yield-related traits. (A) Yield and yield components. Harvest index was calculated as the ratio of the mature seeds dry weight to the above-ground dry weight. Values are the mean and SE of four replicates. Different letters above the bars indicate significant differences (P<0.05), and n.s. indicates not significant differences (p>0.05) by Tukey’s test. (B) Schematic diagram showing the relationships among the yield-related traits. Above DW: above-ground dry weight, DWA seed FW: fresh weight of a mature seed, A seed DW: dry weight of a mature seed, Flower N: cumulative number of open flowers at 49 DAS, HI: harvest index, Leaf DW: leaf dry weight, PMS: percentage of mature seeds, Pod DW: pod dry weight, Pod N: pod number per plant, Pod SR: rate of pod set, Seed DW: seed dry weight, Seed MC: moisture contents of mature seeds, Seed N: total seed number per plant, Seed N / Pod: seed number in a pod, Stem DW: stem dry weight, Yield: seed fresh weight per plant. Pod SR was calculated as the ratio of Pod N to Flower N. Values are the correlation coefficient (n=6, the relationships with Flower N and Pod SR; n=8, the other relationships). *, ** and *** indicates significance at P<0.05, P<0.01 and P<0.001, respectively. Dash lines indicate insignificant relationships, while solid lines indicate significant relationships(P<0.05). The boldness of the lines represents the strength of the correlation, with bolder lines indicating stronger correlation. Red lines represent positive correlation, while blue lines represent negative correlation.

To further investigate yield-determining factors, relationships among traits were examined (Fig. 2B). Total seed number per plant and percentage of mature seeds were significantly positively correlated with fresh seed yield (r = 0.915 and 0.805, respectively), although the percentage of mature seeds did not significantly differ among cultivation systems. Meanwhile, the fresh weight of a mature seed showed a weaker correlation with fresh seed yield (r = 0.479). Among the factors contributing to total seed number, both pod number and seeds per pod were significantly correlated with total seed number (r = 0.994 and 0.930, respectively). Moreover, pod number was more strongly correlated with pod set rate than with flower number (r = 0.974 and 0.624, respectively). Additionally, above-ground dry matter and harvest index were significantly positively correlated with fresh seed yield (r = 0.986 and 0.942, respectively). Above-ground dry matter was also significantly correlated with the dry matter of individual organs. Taken together, these results indicate that pod and seed formation, as well as biomass production, are critical for achieving high yields in plant factories, whereas seed-filling traits such as fresh seed weight and percentage of mature seeds play a lesser role.

### Seed components

The sucrose content was significantly higher in NFT and ROC than in MIST for both cultivars (Fig. 3A). Additionally, in Sayane, NFT and ROC had sucrose levels comparable to FIELD. In contrast, fructose and glucose contents were highest in MIST, although glucose levels did not show statistically significant differences for either cultivar. Total soluble sugars content followed a similar trend to sucrose, which was the most abundant sugar. Starch content was higher in NFT and ROC than in MIST and FIELD for Sayane, whereas for Ooyukimidori, starch accumulation was highest in NFT among the plant factory systems and was comparable to FIELD.

**Fig. 3.**
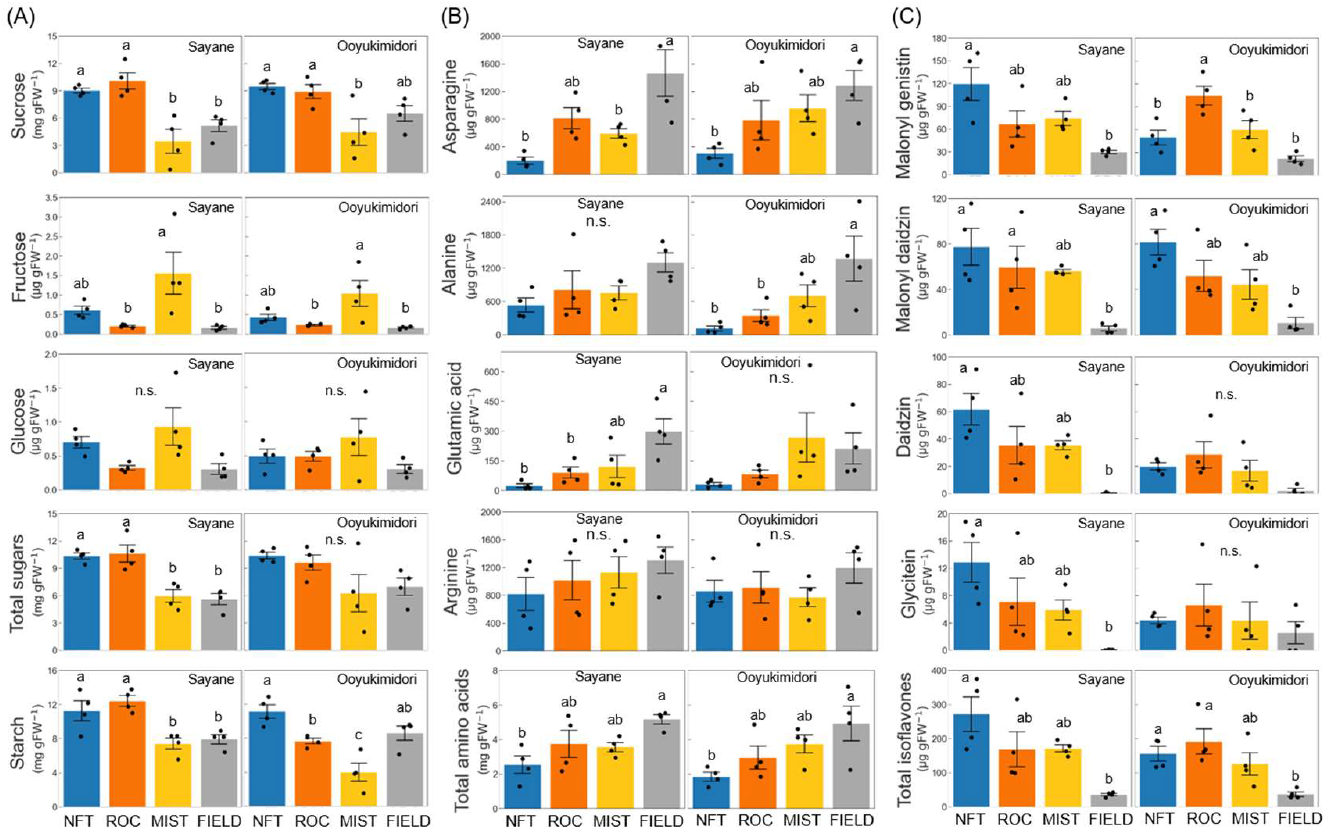
Components in mature seeds. (A) Soluble sugars and starch content. (B) Free amino acids content. (C) Isoflavones content. Total sugars content was calculated as the sum of sucrose, fructose and glucose content. Total free amino acids content was calculated as the sum of amino acids detected in HPLC. Total isoflavones content was calculated as the sum of malonyl genistin, malonyl daidzin, daidzin, and glycitein. Values are the mean and SE of four replicates. Different letters above the bars indicate significant differences (P<0.05), and n.s. indicates not significant differences (p>0.05) by Tukey’s test.

Among free amino acids, no significant differences were found among the plant factory systems (Figs. 3B, S4). However, NFT had significantly lower levels than FIELD in specific amino acids, such as asparagine (both cultivars), alanine (Ooyukimidori), and glutamic acid (Sayane). Total free amino acid content also did not significantly differ among the plant factory systems, but NFT showed significantly lower levels compared with FIELD for both cultivars.

The levels of isoflavones, key functional components in edamame, were also analyzed, including malonyl genistin, malonyl daidzin, daidzin, and glycitein (Fig. 3C). For Sayane, NFT tended to accumulate higher levels of all detected isoflavones, although no significant differences were observed among plant factory systems. For Ooyukimidori, ROC had significantly higher malonyl genistin levels than NFT and MIST, while NFT had moderately higher malonyl daidzin levels. Total isoflavone content was significantly higher in NFT for Sayane and in NFT and ROC for Ooyukimidori compared with FIELD.

### Principal component analysis

To comprehensively evaluate each cultivation system in terms of yield, eating quality, and nutritional properties, principal component analysis (PCA) was performed using multiple parameters, including fresh seed yield, total seed number, harvest index, above-ground dry weight, total amino acids content, total soluble sugars content, and total isoflavones content (Fig. 4). The first two principal components, PC1 and PC2, accounted for 71.1% and 21.8% of the total variance, respectively. PC1 was positively associated with fresh seed yield, total seed number, above-ground dry weight, harvest index, and total soluble sugars content. PC2 was positively associated with total amino acids content and negatively associated with total isoflavones content. NFT had the highest PC1 score and a moderately lower PC2 score for both cultivars. ROC had a slightly higher PC1 score than MIST but remained lower than NFT. Additionally, ROC showed a slightly positive PC2 score for Sayane and a slightly negative PC2 score for Ooyukimidori. MIST had the lowest PC1 and PC2 scores for both cultivars. FIELD had a slightly lower PC1 score, but its PC2 score was higher than that of the plant factory cultivation systems. These results demonstrate that NFT was the most suitable cultivation system for edamame in a plant factory, followed by ROC and MIST. However, NFT-grown edamame exhibited significantly lower total amino acid content than field-grown counterparts.

**Fig. 4.**
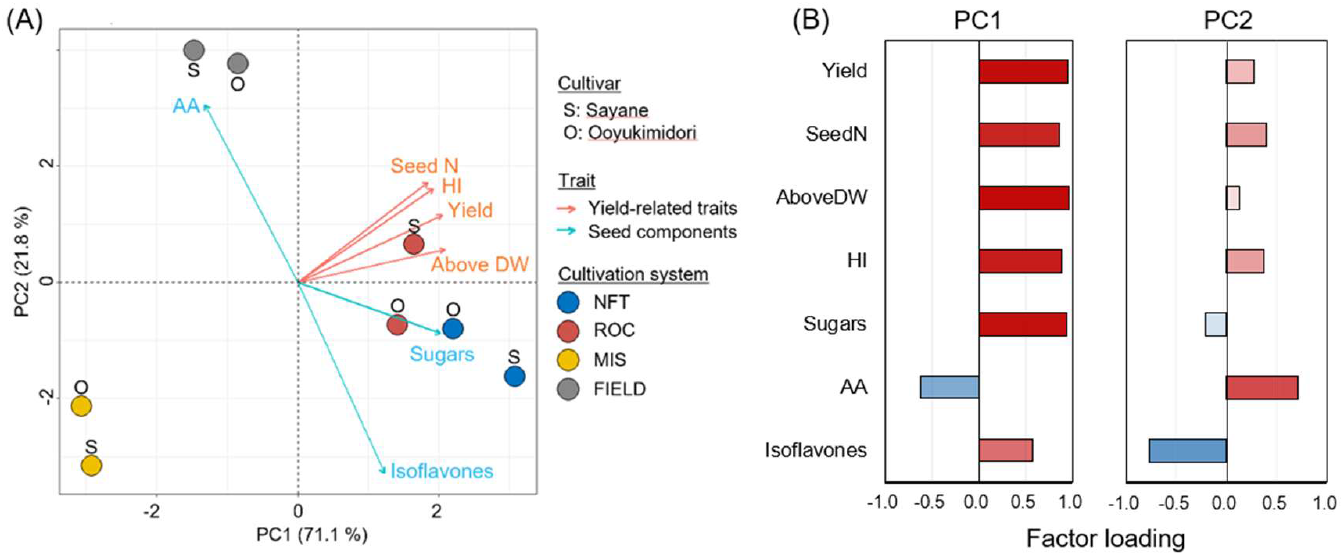
Principal component analysis (PCA) of yield-related traits and seed components. (A) PCA biplot of the cultivation systems with the selected traits, (B) Factor loadings of each trait. AA: total free amino acids content, AboveDW: above-ground dry weight, SeedN: seed number per plant, HI: harvest index, Isoflavones: total isoflavones contents, Sugars: total soluble sugar content, Yield: seed fresh weight per plant. The first two principal components, PC1 and PC2, accounted for 71.1% and 21.8% of the total variance, respectively. Arrows indicate the strength of the trait’s effect on the first two PCs.

## Discussion

A plant factory is a promising model for sustainable and efficient food production in limited spaces. However, the range of crops suitable for plant factories remains restricted, with most systems optimized for leafy vegetables due to their short growth cycles and simpler environmental requirements. Expanding plant factory applications to include fruiting vegetables and protein-rich crops is crucial for diversifying global food production and ensuring future food security. Edamame is a nutrient-rich, high-protein legume with growing global demand^16,17^, yet its hydroponic cultivation remains largely unexplored. In this study, we assessed the feasibility of edamame cultivation in plant factories by comparing three hydroponic systems—NFT, ROC, and MIST—which differ in nutrient delivery and substrate properties (Fig. 1). Our results demonstrated that NFT hydroponics was the most effective system, achieving high yields and superior eating quality comparable to or exceeding conventional field cultivation. These findings establish NFT as a viable method for fruiting vegetable production in controlled environments, paving the way for broader crop diversification in plant factories.

### Yield Optimization Through Key Agronomic Traits

Achieving high fresh seed yield requires understanding the key factors influencing productivity. Our analysis revealed that pod number was the primary determinant of yield, with pod set rate playing a larger role than flower number (Fig. 2B). Additionally, seed number per pod contributed significantly to total seed count, while seed-filling traits, such as individual seed weight and percentage of mature seeds, had minimal impact. These results indicate that increasing both pod number and seed number per pod is crucial for maximizing yield in edamame cultivation. Moreover, we observed a strong correlation between fresh seed yield, above-ground biomass, and harvest index (Fig. 2B). While a higher harvest index reflects greater efficiency in producing harvestable parts, achieving this balance in plant factories—where minimizing plant residues is essential^30^—is challenging. Given that biomass accumulation increases with higher nutrient availability in controlled environments^31^, optimizing the harvest index while sustaining high yields requires further advances in cultivar selection, nutrient management, and environmental control strategies.

### Enhancing Eating Quality and Nutritional Value

Edamame’s taste and nutritional quality are largely determined by seed composition, particularly soluble sugars and amino acids^22,23^. Genotypes with higher levels of sucrose, aspartic acid, glutamic acid, and alanine are associated with better taste^20^. In our study, we observed significant variations in sugar content across cultivation systems, with NFT and ROC accumulating higher sucrose levels than MIST (Fig. 3A). Since sucrose is the predominant sugar in edamame^32^, the increased sucrose levels in NFT and ROC indicate more active sugar and starch metabolism, contributing to enhanced sweetness and greater consumer appeal (Figs. 3A, 5). Additionally, NFT and ROC exhibited significantly higher sucrose levels than FIELD, further reinforcing their potential for high-quality edamame production (Figs. 6, S5).

**Fig. 5.**
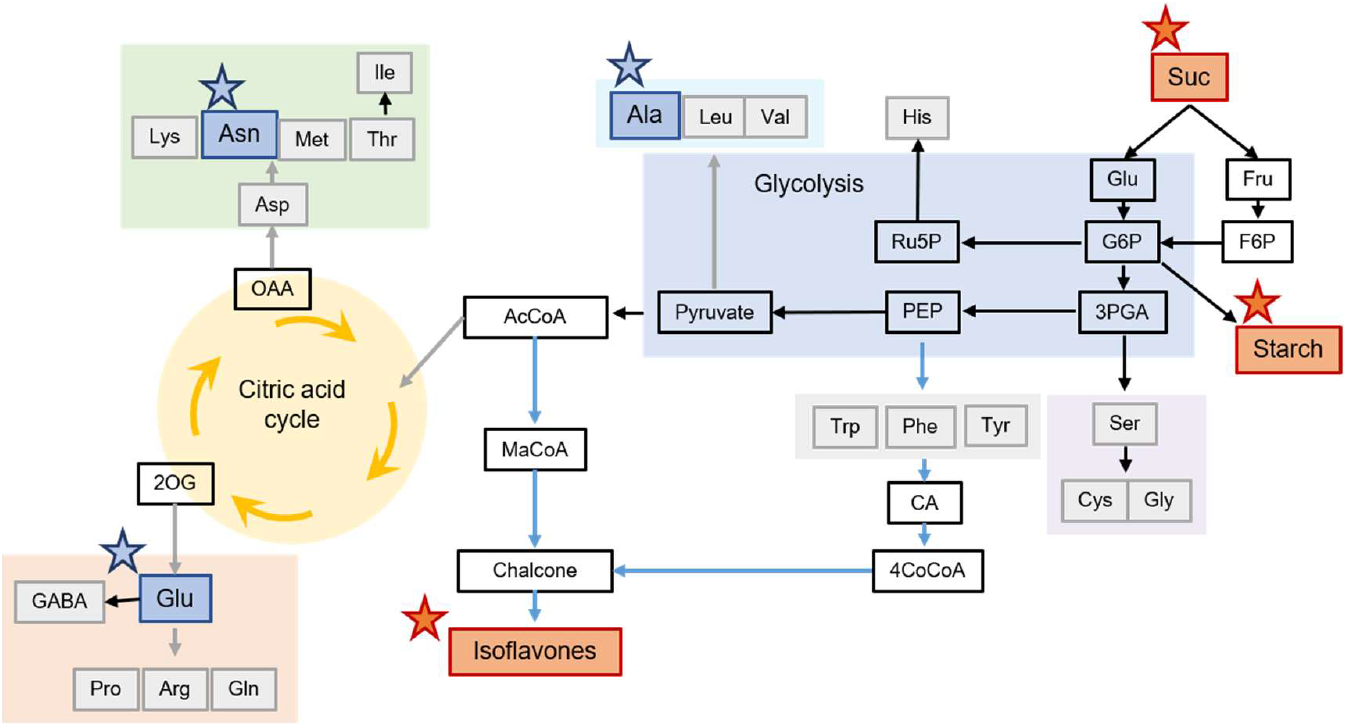
Metabolic map of mature seeds in NFT relative to FIELD. Red star indicates higher abundance and blue star indicate lower abundance in NFT relative to FIELD. AcCoA: acetyl coenzyme A, CA: cinnamic acid, F6P: fructose 6-phosphate, G6P: glucose 6-phosphate, MaCoA: malonyl coenzyme A, PEP: phosphoenolpyruvate, OAA: oxaloacetic acid, Ru5P: ribulose 5-phosphate, 2OG: 2-oxoglutarate, 3PGA: 3-Phosphoglyceric acid, 4CoCoA: 4-coumaroyl coenzyme A.

**Fig. 6.**
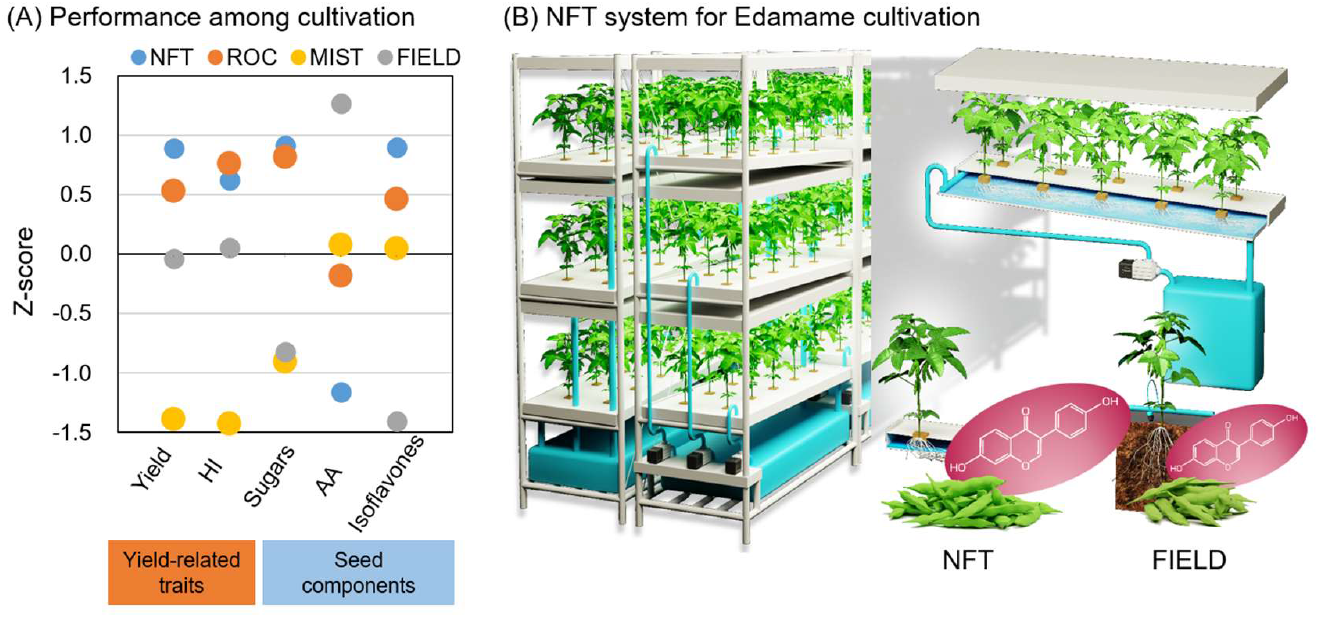
Optimal system for edamame cultivation in an artificial light-type plant factory. (A) Comparison of performance among cultivation systems. AA: total free amino acids content, HI: harvest index, Isoflavones: total isoflavones contents, Sugars: total soluble sugar content, Yield: seed fresh weight per plant. Values indicate standardized z-scores. (B) Conceptual diagram of the introduction of NFT into a plant factory.

We also analyzed isoflavones, which contribute to functional health benefits. While isoflavone content did not differ significantly among plant factory cultivation systems, NFT resulted in higher total isoflavone accumulation than FIELD for both cultivars (Fig. 3C). Given that light quality influences isoflavone biosynthesis in plants^33–35^, our findings suggest that LED lighting promotes isoflavone production more effectively than natural sunlight. However, we observed a negative correlation between isoflavone content and amino acid levels (Fig. S6), indicating a trade-off between these two components. Isoflavones are synthesized via the phenylpropanoid pathway, which originates from phenylalanine, a precursor for multiple secondary metabolites^36^. As isoflavone levels increased, phenylalanine accumulation rose, while major amino acids such as aspartic acid, alanine, and glutamic acid decreased (Figs. 3B, 3C, 5, S4), suggesting that isoflavone biosynthesis competes with amino acid synthesis, leading to reduced free amino acid content. Despite this trade-off, we observed an interaction between cultivar selection and cultivation system for malonyl genistin, the most abundant isoflavone, while no such interaction was observed for major amino acids (Supplemental Table S1). This suggests that strategic breeding could enhance isoflavone content without negatively affecting amino acid levels, offering a promising pathway for further nutritional optimization. Furthermore, amino acids and isoflavones were not correlated with yield (Fig. S7), suggesting that both nutritional quality and yield could be improved simultaneously.

### NFT Hydroponics: The Optimal System for Edamame Cultivation

A comprehensive evaluation of the cultivation systems using principal component analysis (PCA) (Fig. 4) incorporated yield, eating quality, and nutritional attributes. PC1 was primarily associated with yield-related traits and total sugar content, and NFT exhibited the highest PC1 score, confirming it as the most effective system for optimizing both productivity and quality (Figs. 6, S5). While NFT has traditionally been used for leafy vegetables, studies have suggested its potential for fruiting vegetables such as tomatoes, due to its high water-use efficiency and ability to support high-yield production^37^. Our study confirms that NFT is also well-suited for edamame, demonstrating that fruiting vegetables can be successfully cultivated in controlled environments with optimized hydroponic systems. Another key advantage of NFT is its potential for multi-layered cultivation, significantly enhancing space utilization and production efficiency in plant factories. Since early pinching does not negatively impact edamame yield^38^, implementing strategic growth control techniques could further increase vertical farming efficiency, making NFT a scalable and economically viable solution for high-density plant factory production.

### Future Perspectives: From Urban Agriculture to Space Farming

The successful cultivation of high-protein crops like edamame in plant factories has profound implications for global food security, urban farming, and space agriculture. As population growth, climate change, and resource constraints drive demand for efficient and sustainable food production systems, controlled environment agriculture must expand beyond leafy greens to include nutrient-rich legumes and fruiting vegetables. Furthermore, as space exploration advances, long-duration missions and extraterrestrial settlements will require self-sufficient, resource-efficient food systems^29^. While lettuce and other leafy vegetables have been successfully cultivated in space, a nutritionally diverse diet necessitates the inclusion of fruiting vegetables and protein-rich crops. Edamame, with its high protein content, essential amino acids, and functional isoflavones, is an ideal candidate for future space farming. Its successful NFT-based cultivation under artificial lighting (Figs. 6, S5) suggests that it could be adapted for controlled environments beyond Earth, ensuring a balanced, nutrient-dense food supply for astronauts. By leveraging advanced hydroponic systems, fruiting vegetables like edamame could become a cornerstone of sustainable food production for long-duration space missions and extraterrestrial agriculture.

## Conclusion

The present study establishes NFT hydroponics as the optimal system for edamame cultivation in plant factories, achieving higher yields, superior eating quality, and enhanced nutritional properties compared to other hydroponic methods (Figs. 6, S5). Our findings provide strong evidence that fruiting vegetables can be successfully integrated into controlled environment agriculture, expanding the scope of plant factories beyond leafy greens. As urban and space agriculture evolve, the successful integration of high-protein crops like edamame will be essential for enhancing food security, improving resource efficiency, and supporting long-term agricultural sustainability. This study marks a significant step forward in controlled environment agriculture, paving the way for expanded crop diversity and technological advancements in terrestrial and extraterrestrial food production.

## Materials and methods

### Plant materials and cultivation systems

Two edamame cultivars, Sayane (Snow Brand Seed Co., Ltd., Hokkaido, Japan) and Ooyukimidori (Otani Seed Co., Ltd., Hokkaido, Japan), were used in this study. Seeds were sown in rockwool cube sheets (36 mm × 36 mm × 39 mm per cube) placed in bottom-watering trays (318 mm × 634 mm × 58 mm) filled with pure water and covered with a blackout curtain. The blackout curtain was removed three days after sowing (DAS), and germinated seedlings were grown under a light intensity of 530 ± 12 µmol m^−2^ s^−1^ for four days until primary leaves emerged. Seedlings were then transferred to a nutrient solution (EC = 0.5 dS m^−1^) prepared using OAT House No.1/No.2 (OAT Agrio Co., Ltd., Tokyo, Japan) and grown for an additional seven days. At 14 DAS, seedlings with fully expanded primary leaves were selected and transplanted into each cultivation system.

Experiments were conducted in a plant factory, where plant growth shelves (L1350 mm × W380 mm × H1880 mm) and culture solution tanks (L785 mm × W370 mm × H325 mm) were used. White reflective films were placed on both sides of the shelves to enhance light intensity. The light intensity at the canopy level was maintained at 520 ± 17 µmol m^−2^ s^−1^ across all cultivation systems by adjusting the light source intensity and the distance between plants and light sources. The nutrient solution EC was set at 0.5 dS m^−1^ on the first day after transplanting to minimize environmental stress and adjusted to 1.5 ± 0.05 dS m^−1^ from the second day onward. EC adjustments were performed twice a week, and the nutrient solution was fully replaced once during the mid-growth stage. Three hydroponic cultivation systems—nutrient film technique (NFT), rockwool culture (ROC), and mist culture (MIST)—were examined in this study. Details of each system are described below.

### 1) Nutrient Film Technique (NFT)

NFT cultivation was performed on multi-layered plant growth shelves (Fig. S8). Each cultivation layer measured L1350 mm × W380 mm and was equipped with a growing bed (L1040 mm × W295 mm × H35 mm) and a root-support panel (L600 mm × W295 mm). Each layer accommodated five plants per cultivar. The culture solution was supplied from a container tank positioned at the base of the shelf and flowed continuously through the bottom 10 mm of the root zone at a flow rate of 1.08 ± 0.12 m^3^ h^−1^. The effluent solution was recirculated back into the container tank.

### 2) Rockwool Culture (ROC)

ROC cultivation was also conducted on multi-layered plant growth shelves (Fig. S9). Each layer measured L1350 mm × W380 mm and contained a rockwool slab (Grodan Vital 4716M, L900 mm × W195 mm × H75 mm) as a growth medium. Five plants per cultivar were transplanted onto the rockwool slabs. Drip irrigation was provided via plug-in sprinklers (GKS101, Takagi Co., Ltd., Fukuoka, Japan), which were placed at the base of each plant and at both ends of the rockwool slabs. Irrigation was conducted once daily for 5 minutes at the midpoint of the photoperiod to prevent excessive increases in rockwool temperature. The irrigation volume was optimized to saturate the rockwool slab without excess pooling of the culture solution. Any surplus solution was drained from the back of the rockwool slab and collected in the container tank for reuse.

### 3) Mist Culture (MIST)

MIST cultivation was performed in containers (L785 mm × W370 mm × H325 mm) (Fig. S10), with five plants per cultivar. Plants were secured in planting holes on polystyrene foam using half-sized plug tray holes (128-hole tray, Tomikawa Chemical Industry Co., Ltd., Aichi, Japan). An ultrasonic mist generator (Mini-Mist Maker HS0276, AGPTTEK) was placed inside the container to generate mist for 20 seconds every 40 seconds, ensuring full mist coverage without excessive temperature increases. The culture medium depth was maintained at 100 mm. A circulating pump (Aheim Compacton 1000, Kamihata Fish Industry Group, Hyogo, Japan) and an aeration device (E-Air 4000 WB, JECS Co., Ltd., Osaka, Japan) were installed to facilitate nutrient circulation and maintain oxygen supply. The lower 10 cm of the root system was immersed in the culture medium, while the upper root zone remained exposed to mist.

### 4) Field Cultivation

For comparison, field-grown plants were cultivated at the Institute for Sustainable Agro-ecosystem Services (ISAS), The University of Tokyo (35°44’N, 139°32’E, Tokyo, Japan) from June 7 to August 24, 2023 (Fig. S11). A compound fertilizer (N: P: K = 3:3:10) was applied as a basal dressing at 200 g m^−2^. Seeds were sown in mulched soil with 100 mm diameter holes, with three seeds per hole sown at a depth of 20 mm. At 7 DAS, seedlings were thinned to one plant per hill.

The planting density was standardized at 12.5 plants m^−2^ (140 mm spacing between plants) across all cultivation systems, including the plant factory and field conditions. To control plant elongation, the growing point was removed at the seventh node using scissors once the seventh node became visible (Fig. S12). Until harvest, thinned leaves were collected to measure dry weight. Plants were harvested when all pods on a plant had fully swollen to their maximum size. The harvest timing was 79 DAS for both plant factory and field-grown crops.

### Environmental condition

Environmental data for the plant factory were recorded using a data logger (TR-76Ui, T&D Corporation, Nagano, Japan), while meteorological data for the field experiment were obtained from a fixed-point meteorological observation device placed in the field. The air temperature in the plant factory remained constant at 26.7°C, whereas in the field, temperatures fluctuated, with a daily mean of 23.5°C, a maximum of 32.1°C, and a minimum of 13.2°C throughout the growing period (Fig. S13A, B). Root zone temperature was monitored using a data logger (TR-52i, T&D Corporation, Nagano, Japan). Root zone temperatures remained stable in NFT and MIST, while in ROC, temperatures increased with LED irradiation, temporarily decreased during drip irrigation, and declined further after lights were turned off (Fig. S13C).

### Measurement of growth traits

The main stem length, leaflet number with a diameter of over 30 mm, and number of nodes with at least one leaflet of over 30 mm on main stem were measured at 16, 20, 22, and 28 days after sowing (DAS). The flower number was counted at 28, 30, 33, 37, 41, 45, and 49 DAS. At maturity stage, plants were harvested and divided into stems, leaves, and pods. Stems and leaves were dried to a constant weight at 80 °C for seven days and weighed for dry matter. Pods were divided according to the seed number in a pod and number and fresh weight of each pod were measured. Seeds larger than 10 mm in diameter were regarded as mature seeds, and the others were regarded as immature seeds. Seed number, fresh weight, and dry weight in each size group were measured.

### Quantification of soluble sugars and starch

At maturity stage, mature seeds were harvested from four plants and stored at −30°C until analyzing seed components. A frozen mature seed was ground to a fine powder using a mortar and pestle. Approximately 50 mg of ground sample was used for measuring soluble sugars and starch contents. Extraction and measuring for sugars and starch were conducted based on the methods of Okamura *et al*.^39^ using glucoamylase (Toyobo Co., Ltd., Osaka, Japan), an F-kit #716260 (J.K. International Co., Ltd., Tokyo, Japan), and a microplate reader, Biotek Synergy H1 (Agilent Technologies Inc., California, US).

### Quantification of free amino acids and isoflavones

Three frozen mature seeds were ground using a mortar and pestle. Approximately 2 g of ground sample was used for measuring free amino acids and isoflavones contents. 500 µL of tissue lysate was mixed with 1.5 mL 99.5% ethanol (EtOH) and extracted with shaking for 5 min at room temperature. After centrifuge, 1.0 mL supernatant was collected. 500 µL of the collected supernatant was used for isoflavones measurement. The remaining supernatant was vacuum dried for 2 h at 40 °C. The dried sample was suspended with 40 µL of EWT solution (EtOH: H2O: triethylamine = 2: 2: 1) and vacuum dried for 1 h at 40 °C. Subsequently, the dried sample was resuspended with 40 µL of EWTP solution (EtOH: H2O: triethylamine: phenyl isothiocyanate = 7: 1: 1: 1) and vacuum dried for 1 h at 40 °C, and then mixed with 40% acetonitrile. This mixture was used for amino acids measurement.

Free amino acids and isoflavones contents were analyzed using high-performance liquid chromatography on a TSKgel Super-ODS (Tosoh Bioscience, Tokyo, Japan) and TSKgel ODS-120T (Tosoh Bioscience, Tokyo, Japan), respectively. Free amino acids were eluted with solvent A (0.1 M ammonium acetate: 5 % acetonitrile = 95: 5) and solvent B (acetonitrile: 1% acetic acid = 4: 6) at a flow rate of 1.0 mL min^−1^ at a column temperature of 40 °C and detected at 254 nm. The gradient elution was conducted as follows: 100% A at 0-3 min, 100-25% A and 0-75 % B at 3-7.5 min, 25% A and 75% B at 7.5-15 min. Isoflavones were eluted with same solvent A and B at a flow rate of 1.0 mL min^−1^ at a column temperature of 40 °C and detected at 254 nm. The gradient elution was conducted as follows: 100% A at 0 min, 100-0% A and 0-100 % B at 0-15 min, 100% B at 15 min. The contents for every amino acid and every isoflavone in a sample were quantified by measuring the relative peak area to the standard.

### Statistical analysis

Analysis of variance (ANOVA) was conducted using R-Studio (v.4.0.3). For each cultivar and cultivation system, significance of differences among mean values was analyzed using Tukey’s test (*P*<0.05) with multcomp package. Principal component analysis (PCA) for growth traits and seed components in each cultivation system was performed using R-Studio (v.4.0.3) with FactoMineR and factoextra packages. The correlation coefficient and regression line were calculated using Microsoft Excel. Z-scores were calculated by the scale function in R-studio (v.4.0.3).

## Data availability statement

Supporting data can be requested by contacting the corresponding author.

## Acknowledgments

This work was supported by KAKENHI (18KK0170, 21H02171, and 24H02277 to W.Y.) from the Japan Society for the Promotion of Science (JSPS).

## Author contributions

T.T., Y.W. and W.Y. designed the experiments. T.T. and Y.W. grew the plants, performed the experiments, and analyzed the data. S.W. and T.S. performed quantifications of free amino acids and isoflavones. T.T., Y.W. and W.Y. prepared the manuscript. All the authors have read and approved the final version of this manuscript.

## Figure legends

**Supplemental Figure S1**. Growth traits at vegetative growth stage.

Values are the mean and SE of five replicates. Different letters above the bars indicate significant differences (P<0.05), and n.s. indicates not significant differences (p>0.05) by Tukey’s test.

**Supplemental Figure S2**. Dry matter at maturity.

Above-ground dry weight was calculated as the sum of the dry weights of stem, leaf, pod, and seed. Values are the mean and SE of four replicates. Different letters above the bars indicate significant differences (P<0.05), and n.s. indicates not significant differences (p>0.05) by Tukey’s test.

**Supplemental Figure S3**. Changes in cumulative number of open flowers.

Values are the mean and SE of five replicates. Different letters above the bars indicate significant differences (P<0.05), and n.s. indicates not significant differences (p>0.05) by Tukey’s test.

**Supplemental Figure S4**. Free amino acids content in mature seeds.

Values are the mean and SE of four replicates. Different letters above the bars indicate significant differences (P<0.05), and n.s. indicates not significant differences (p>0.05) by Tukey’s test.

**Supplemental Figure S5**. Comparison of performance among cultivars in each cultivation system. AA: total free amino acids content, HI: harvest index, Isoflavones: total isoflavones contents, Sugars: total soluble sugar content, Yield: seed fresh weight per plant. Values indicate standardized z-scores.

**Supplemental Figure S6**. Relationship of total isoflavones content with total amino acids content. Values are the mean and SE of four replicates. * indicates significant differences at the 0.05 level.

**Supplemental Figure S7**. Relationships of the fresh seed yield with total amino acids and total. Values are the mean and SE of four replicates.

**Supplemental Figure S8**. Equipment in nutrient film technique (NFT).

(A, B) Cultivation appearance in NFT, (C) Supply of culture medium to root zone, (D) Circulation system of culture medium, (E) A pump used to circulate culture medium.

**Supplemental Figure S9**. Equipment in rockwool culture (ROC).

(A, B) Cultivation appearance in ROC, (C) Drip irrigation at the base of plant using a sprinkler,

(D) A plug-in sprinkler used in ROC.

**Supplemental Figure S10**. Equipment in mist culture (MIST).

(A, B) Cultivation appearance in MIST, (C) Supply of culture medium to root zone by mist filling

(D) An ultrasonic mist generator used in MIST.

**Supplemental Figure S11**. Plant cultivation in the field (FIELD). (A, B) Field cultivation.

**Supplemental Figure S12**. Pinching treatment.

Pinching treatment was conducted by cutting off the growth point, the upper 10mm of the seventh node, with scissors at the time the seventh node was visible. In fact, this treatment was conducted as needed after 28DAS.

**Supplemental Figure S13**. Environmental condition from July 9, 2023 to July 12, 2023 in plant factory and field. (A) Air temperature and relative humidity in field, (B) Air temperature and relative humidity in plant factory, (C) Root zone temperature in each cultivation system of plant factory.

