## Supplementary Material for "Sustainable Edamame Production in an Artificial Light Plant Factory with Improved Yield and Quality"

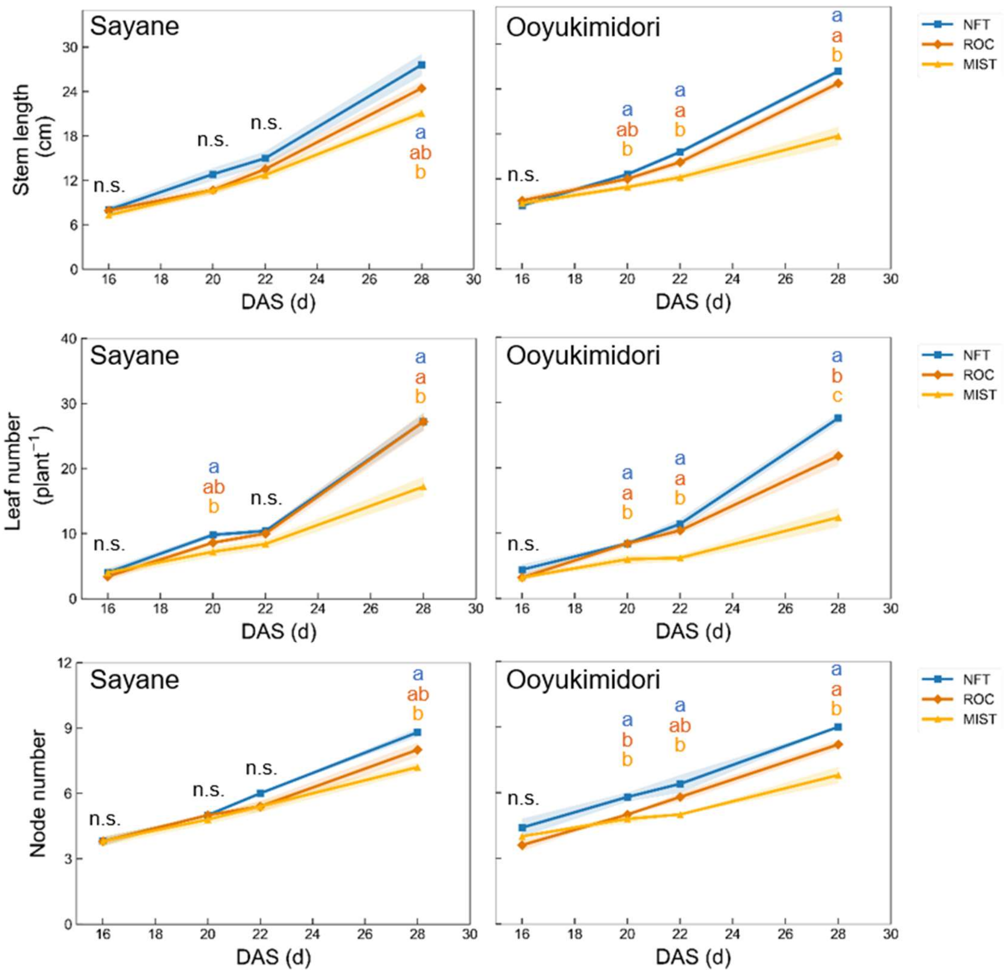


**Supplemental Figure S1.** Growth traits at vegetative growth stage.

Values are the mean and SE of five replicates. Different letters above the bars indicate significant differences (P<0.05), and n.s. indicates not significant differences (p>0.05) by Tukey's test.

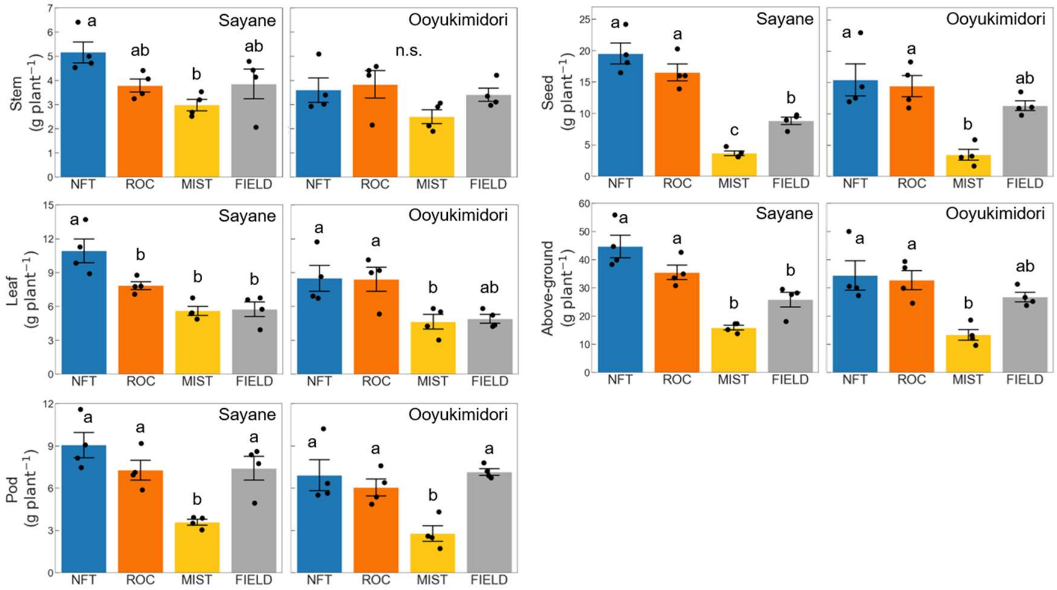


**Supplemental Figure S2.** Dry matter at maturity.

Above-ground dry weight was calculated as the sum of the dry weights of stem, leaf, pod, and seed. Values are the mean and SE of four replicates. Different letters above the bars indicate significant differences ($P<0.05$), and n.s. indicates not significant differences ($p>0.05$) by Tukey's test.

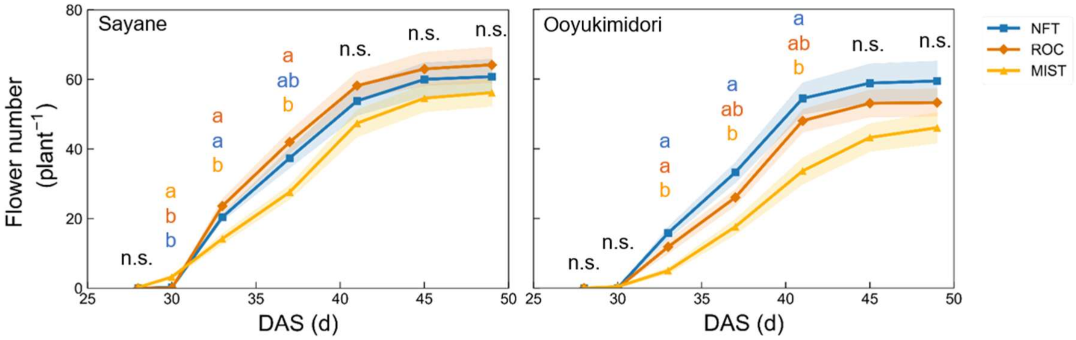


**Supplemental Figure S3.** Changes in cumulative number of open flowers.

Values are the mean and SE of five replicates. Different letters above the bars indicate significant differences (P<0.05), and n.s. indicates not significant differences (p>0.05) by Tukey's test.

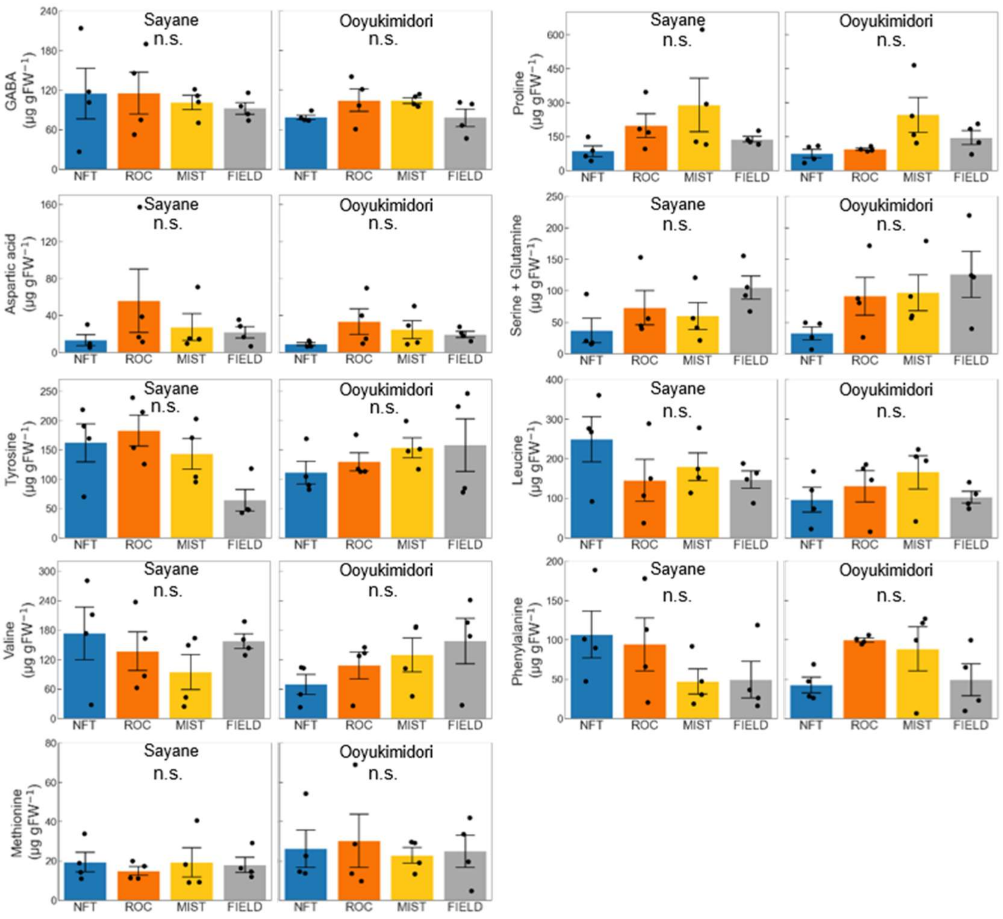


**Supplemental Figure S4.** Free amino acids content in mature seeds.

Values are the mean and SE of four replicates. Different letters above the bars indicate significant differences ($P<0.05$), and n.s. indicates not significant differences ($p>0.05$) by Tukey's test.

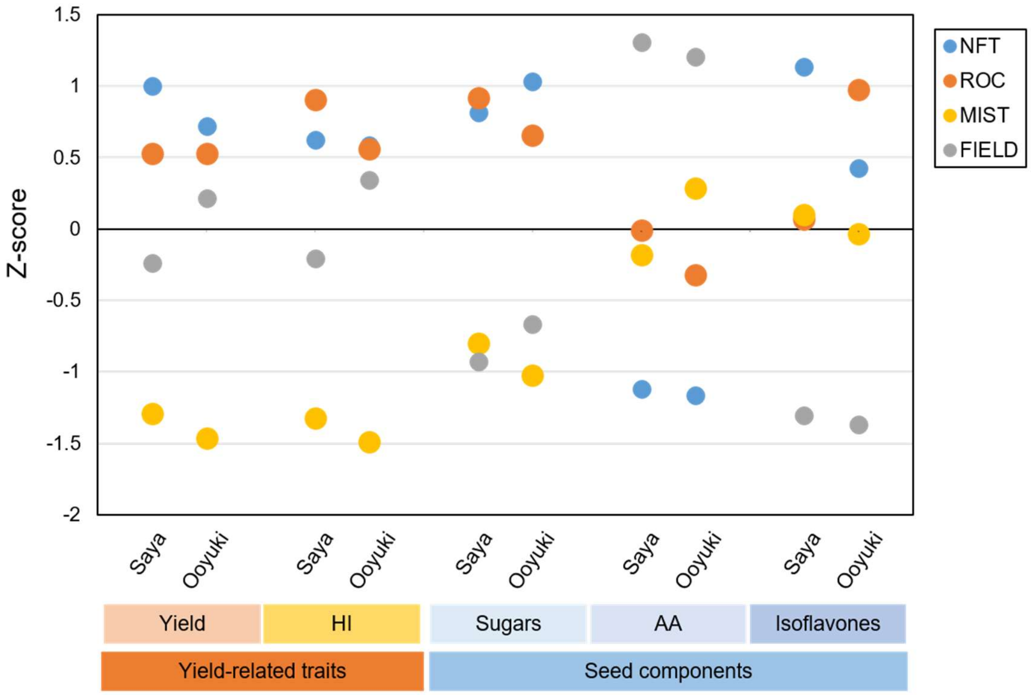


**Supplemental Figure S5.** Comparison of performance among cultivars in each cultivation system. AA: total free amino acids content, HI: harvest index, Isoflavones: total isoflavones contents, Sugars: total soluble sugar content, Yield: seed fresh weight per plant. Values indicate standardized z-scores.

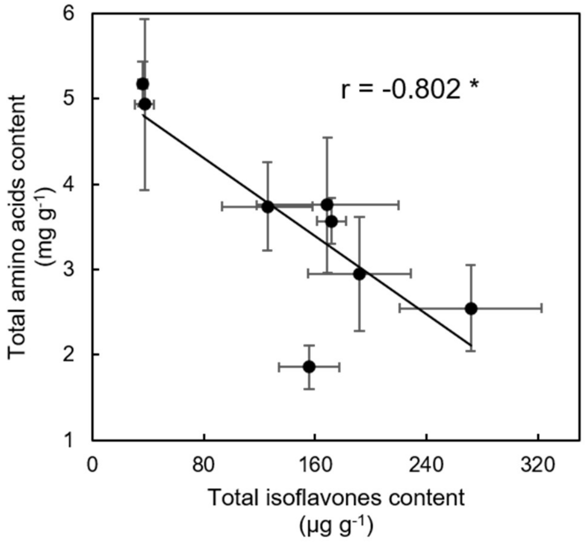


**Supplemental Figure S6.** Relationship of total isoflavones content with total amino acids content.

Values are the mean and SE of four replicates. * indicates significant differences at the 0.05 level.

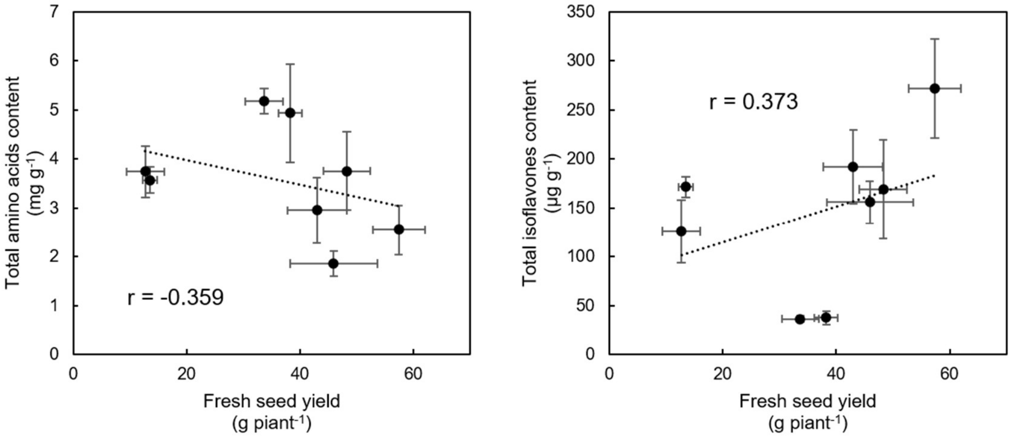


**Supplemental Figure S7.** Relationships of the fresh seed yield with total amino acids and total. Values are the mean and SE of four replicates.

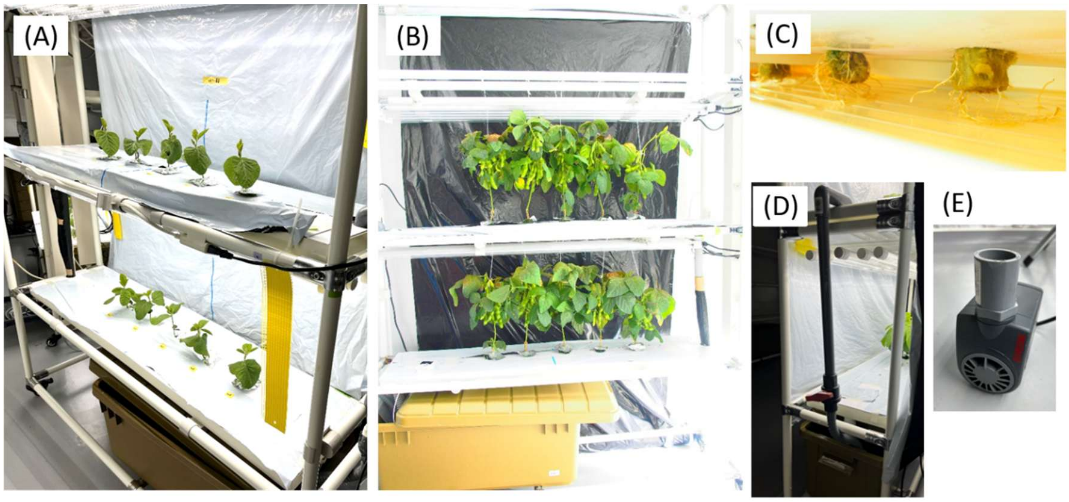


**Supplemental Figure S8.** Equipment in nutrient film technique (NFT).

(A, B) Cultivation appearance in NFT, (C) Supply of culture medium to root zone, (D) Circulation system of culture medium, (E) A pump used to circulate culture medium.

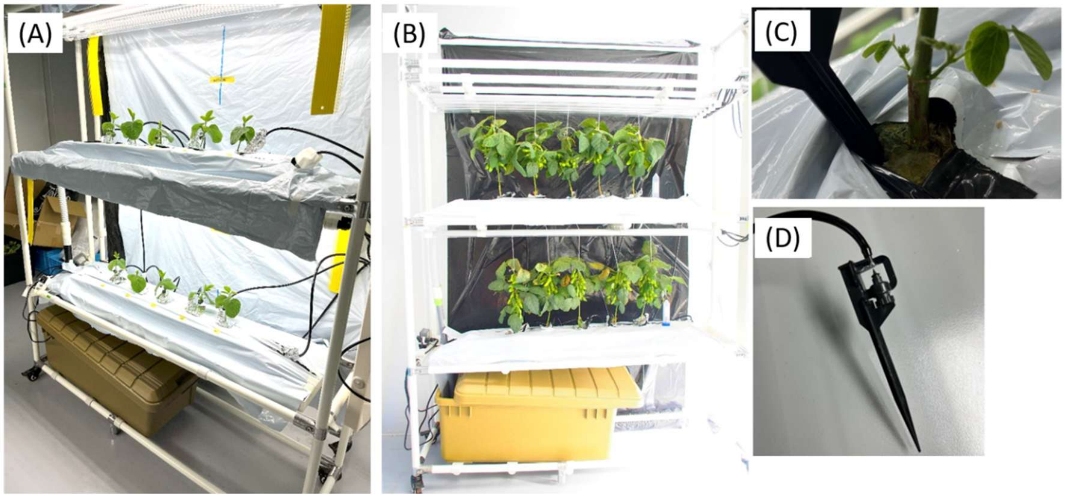


**Supplemental Figure S9.** Equipment in rockwool culture (ROC).

(A, B) Cultivation appearance in ROC, (C) Drip irrigation at the base of plant using a sprinkler, (D) A plug-in sprinkler used in ROC.

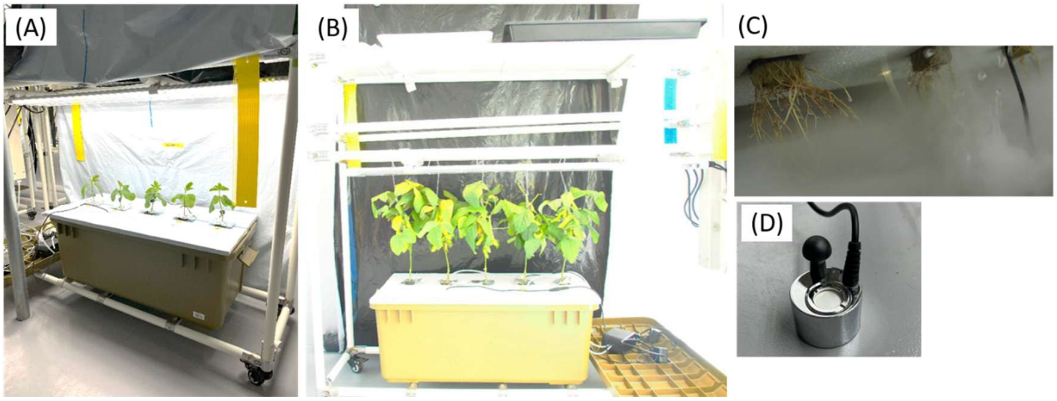


**Supplemental Figure S10.** Equipment in mist culture (MIST).

(A, B) Cultivation appearance in MIST, (C) Supply of culture medium to root zone by mist filling

(D) An ultrasonic mist generator used in MIST.

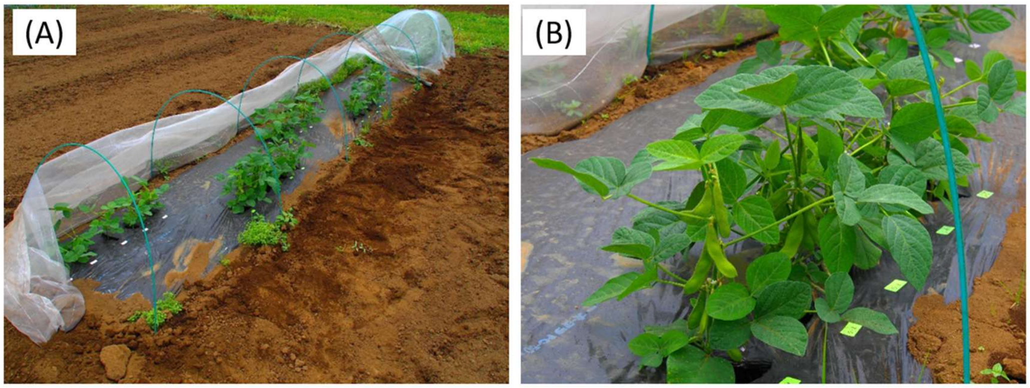


**Supplemental Figure S11.** Plant cultivation in the field (FIELD).

(A, B) Field cultivation.

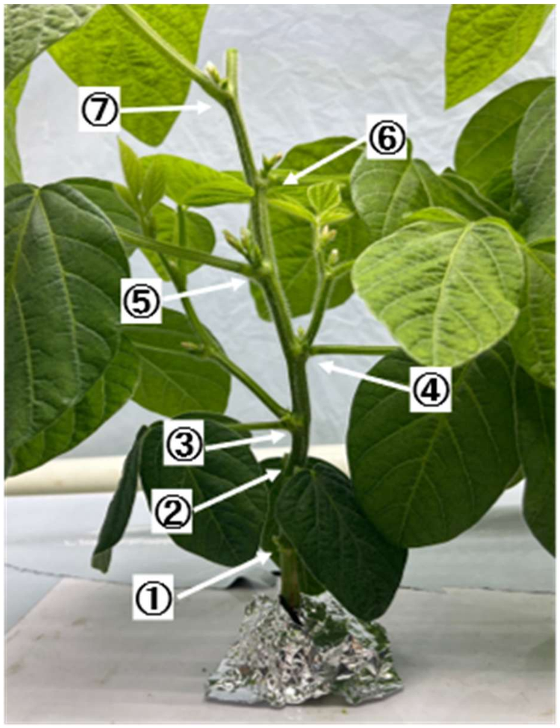


**Supplemental Figure S12.** Pinching treatment.

Pinching treatment was conducted by cutting off the growth point, the upper 10mm of the seventh node, with scissors at the time the seventh node was visible. In fact, this treatment was conducted as needed after 28DAS.

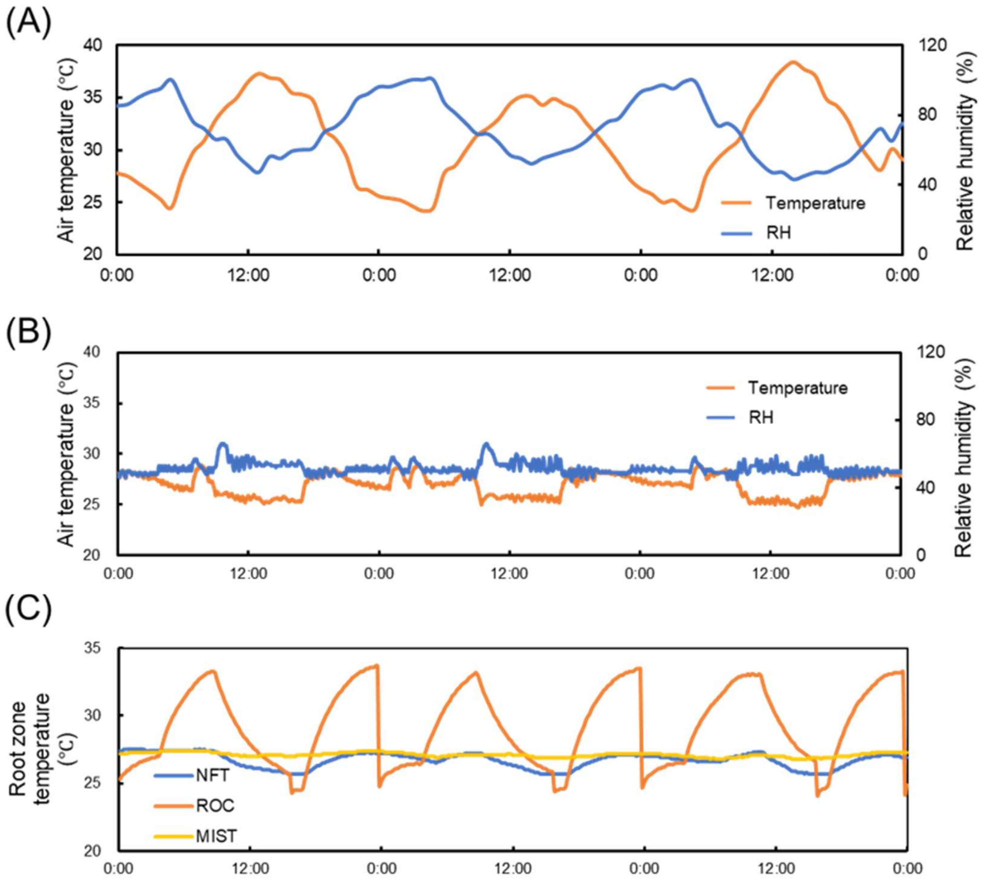


**Supplemental Figure S13.** Environmental condition from July 9, 2023 to July 12, 2023 in plant factory and field. (A) Air temperature and relative humidity in field, (B) Air temperature and relative humidity in plant factory, (C) Root zone temperature in each cultivation system of plant factory.

**Supplemental Table S1.** Analysis of variance (ANOVA) for the contents of isoflavones and free amino acids.

| Cultivar | System | Isoflavones | | | | | Free amino acids | | | | |
|---|---|---|---|---|---|---|---|---|---|---|---|
| | | Malonyl genistin | Malonyl daidzin | Daidzin | Glycitein | Total isoflavones | Asparagine | Alanine | Glutamic acid | Arginine | Total amino acids |
| (C) | (S) | (μg gFW$^{-1}$) | (μg gFW$^{-1}$) | (μg gFW$^{-1}$) | (μg gFW$^{-1}$) | (μg gFW$^{-1}$) | (μg gFW$^{-1}$) | (μg gFW$^{-1}$) | (μg gFW$^{-1}$) | (μg gFW$^{-1}$) | (mg gFW$^{-1}$) |
| Sayane | NFT | 119.5 a | 77.5 a | 61.8 a | 12.9 a | 271.7 a | 201 b | 538 ab | 24 a | 819 a | 2.55 ab |
| | ROC | 66.9 abc | 59.4 ab | 35.4 ab | 7.1 ab | 168.9 ab | 813 ab | 812 ab | 91 a | 1020 a | 3.75 ab |
| | MIST | 74.1 abc | 56.0 ab | 35.4 ab | 5.9 ab | 171.4 ab | 595 ab | 757 ab | 122 a | 1130 a | 3.57 ab |
| | FIELD | 30.0 c | 5.8 b | 0.4 b | 0.1 b | 36.2 b | 1469 a | 1306 a | 299 a | 1305 a | 5.17 a |
| Ooyukimidori | NFT | 49.8 bc | 81.8 a | 19.9 b | 4.4 ab | 155.9 ab | 306 b | 116 b | 33 a | 861 a | 1.86 b |
| | ROC | 104.6 ab | 52.1 ab | 28.5 ab | 6.6 ab | 191.9 a | 783 ab | 348 ab | 84 a | 914 a | 2.95 ab |
| | MIST | 60.1 bc | 44.5 ab | 16.8 b | 4.3 ab | 125.7 ab | 957 ab | 706 ab | 268 a | 772 a | 3.74 ab |
| | FIELD | 22.1 c | 10.6 b | 2.3 b | 2.6 ab | 37.6 b | 1288 a | 1375 a | 213 a | 1195 a | 4.93 a |
| mean | Sayane | 72.6 a | 49.7 a | 33.2 a | 6.5 a | 162.1 a | 770 a | 853 a | 134 a | 1069 a | 3.76 a |
| | Ooyukimidori | 59.2 a | 47.2 a | 16.8 b | 4.5 a | 127.7 a | 834 a | 636 a | 149 a | 935 a | 3.37 a |
| mean | NFT | 84.7 a | 79.6 a | 40.8 a | 8.6 a | 213.8 a | 254 b | 327 b | 29 b | 840 a | 2.20 b |
| | ROC | 85.8 a | 55.8 a | 32.0 a | 6.9 ab | 180.4 a | 798 b | 580 b | 87 ab | 967 a | 3.35 b |
| | MIST | 67.1 a | 50.3 a | 26.1 a | 5.1 ab | 148.6 a | 776 b | 731 ab | 195 ab | 951 a | 3.65 ab |
| | FIELD | 26.1 b | 8.2 b | 1.3 b | 1.3 b | 36.9 b | 1379 a | 1340 ab | 256 a | 1250 a | 5.05 a |
| ANOVA | Cultivar (C) | n.s. | n.s. | ** | n.s. | n.s. | n.s. | n.s. | n.s. | n.s. | n.s. |
| | System (S) | *** | *** | *** | * | *** | *** | ** | ** | n.s. | *** |
| | C×S | ** | n.s. | n.s. | n.s. | n.s. | n.s. | n.s. | n.s. | n.s. | n.s. |

Values are the mean and SE of four replicates. Different letters indicate significant differences at the 0.05 level. ** and *** indicate significant differences at the 0.01 and 0.001 levels, respectively, and n.s. indicates not significant differences.
